# A study of machine learning techniques for Automated Karyotyping System

**DOI:** 10.1101/2023.11.16.567473

**Authors:** Kamalpreet Kaur, Renu Dhir

## Abstract

Genetic abnormalities constitute a considerable share of all the existing societal healthcare issues. There has been a dire need for the automation of chromosomal analysis, hence supporting laboratory workers in effective classification and identifying such abnormalities. Nevertheless, with many modern image processing techniques, like Karyotyping, improved the life expectancy and the quality of life of such cases. The standard image-based analysis procedures include Pre-processing, Segmentation, Feature extraction, and Classification of images. When explicitly considering Karyotyping, the processes of Segmentation and Classification of chromosomes have been the most complex, with much existing literature focusing on the same. Various model-based machine learning models have proven to be highly effective in solving existing issues and building an artificial intelligence-based, autonomous-centric karyotyping system. An autonomous Karyotyping System will connect the pre-processing, Segmentation, and classification of metaphase images. The review focuses on machine learning-based algorithms for efficient classification accuracy. The study has the sole motive of moving towards an effective classification method for karyotype metaphase images, which will eventually predict the fetus’s abnormalities more effectively. The study’s results shall benefit future researchers working in this area.

## 1. Introduction

Population demands improvement in the numeral of genetic diseases with appropriate planning and strategies for better forecasting potential future. A medical ailment known as a congenital disease arises from a genetic mutation or anomaly in an individual’s DNA. The changes in gene structure result in disorders that might be inherited from parents or result from DNA mutation.

*Genetic diseases* can manifest early in infancy or later in life, impacting numerous aspects of a person’s health. A person’s metabolism, general well-being, and physical and mental development are just a few of the areas of health that genetic abnormalities can impact. Thousands of genetic illnesses exist, each with a unique set of symptoms.

Genetic disorder is categorized into three types: Single-gene disorder, Chromosomal disorder and Complex disorder, as shown in Figure 1. An abnormal chromosome can lead to many health issues within the body. The cause of abnormal chromosomal has been the mistake that occurred during the cell division. One or more of these frequently causes chromosomal abnormalities, as shown in Figure 2.

**Figure 1:**
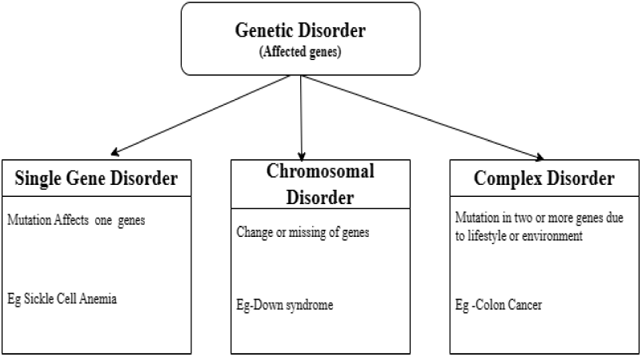
Different levels of chromosomal organization

**Figure 2:**
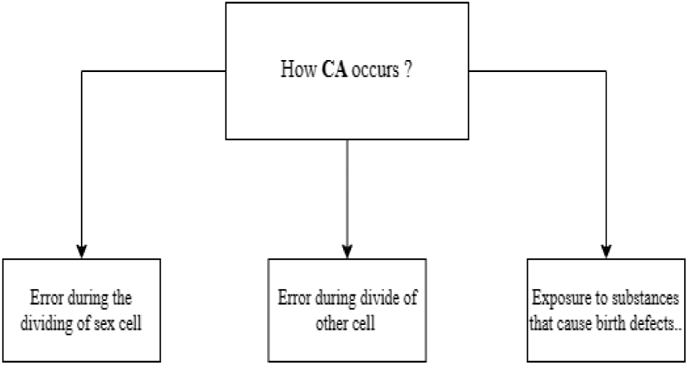
Chromosome abnormalities (C.A.) occurrence.

Every chromosome in our body has a unique structure. Every cell’s nucleus containing chromosomes comprises a thread-like structure of DNA (Deoxyribonucleic acid molecules), protein, and chromatin, shown in Figure 3 [1].

**Figure 3:**
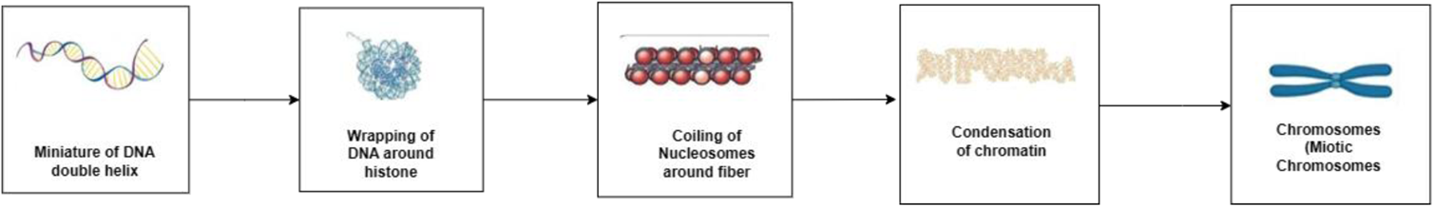
Mitotic chromosomes generated from Deoxyribonucleic acid molecules (DNA)

Chromosome’s physical state and spindle, five distinct phases of the mitosis stage, can be comprehended. These stages are *prophase, prometaphase, metaphase, anaphase, and telophase, as* shown in Figure 4 [1].

**Figure 4:**
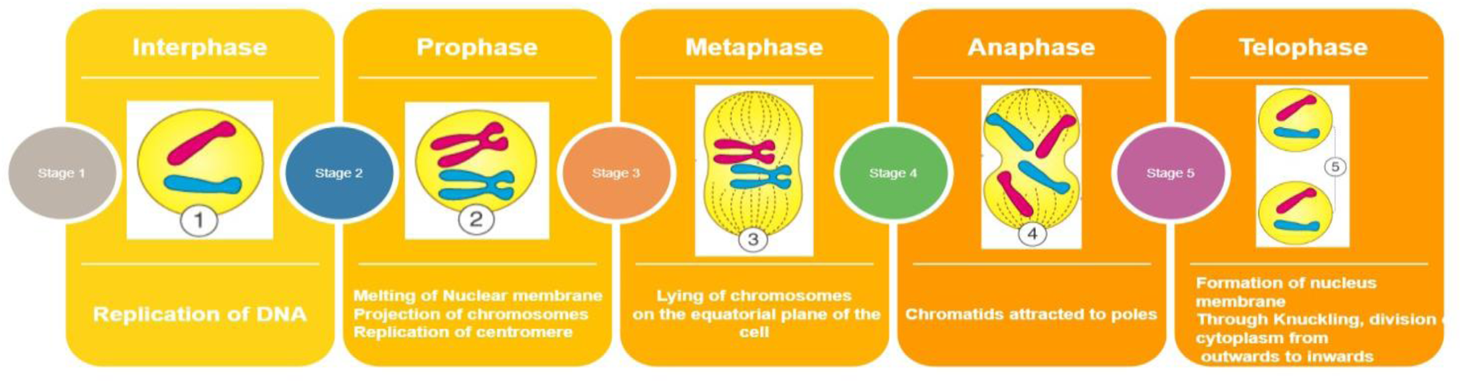
Stages of Mitosis

The cytoplasm division occurs in this phase [2]. Metaphase chromosomes are divided into 24 groups called karyograms using the usual karyotyping method. Humans fetch one set from any parent, obtaining an average pair of 23 chromosomes. In the Karyogram, the first 22 pairs of chromosomes, known as autosomes, facilitate various tasks in human cells. The 23rd pair of sex chromosomes, XX and X.Y., can be found in both forms in all healthy human cells. Chromosomes profoundly impact human health, and any numerical or structural abnormalities in these cells can cause various illnesses, including intellectual deficiencies, congenital deformities, infertility, sexual variations, recurrent pregnancy loss, and even cancer [3]. There are multiple steps in the karyotyping process: *sample collection, chromosome staining, chromosome investigation and cell culture factor.* The scale of karyotyping has been spreading its wings in different domains. *Prenatal Diagnostics* to identify chromosomal abnormalities in fetuses. *Tumor Diagnosis* performs chromosomal abnormalities and genetic alterations linked to specific forms of cancer. *Chromosomal Syndromes are* essential in diagnosing chromosomally aberrant genetic diseases such as Turner syndrome, Down syndrome, and Klinefelter syndrome. An automatic karyotyping system is a specialized computer-based technology or software that helps analyze and interpret karyotypes, whether they belong to humans or other species. A karyotype is a picture of a person’s chromosomes, usually ordered by size and banding patterns, usually in pairs. It is a crucial approach for detecting and investigating chromosomal abnormalities, including structural chromosomal rearrangements and aneuploidy (abnormal chromosome number), in cytogenetics, genetics, and clinical diagnostics. The computer-aided karyotyping system is completed in various stages, such as Image Capture, Image Analysis, Pairing and Sorting, Classification, Aberration Detection, Reporting and Database Integration, as shown in Figure 5.

**Figure 5:**
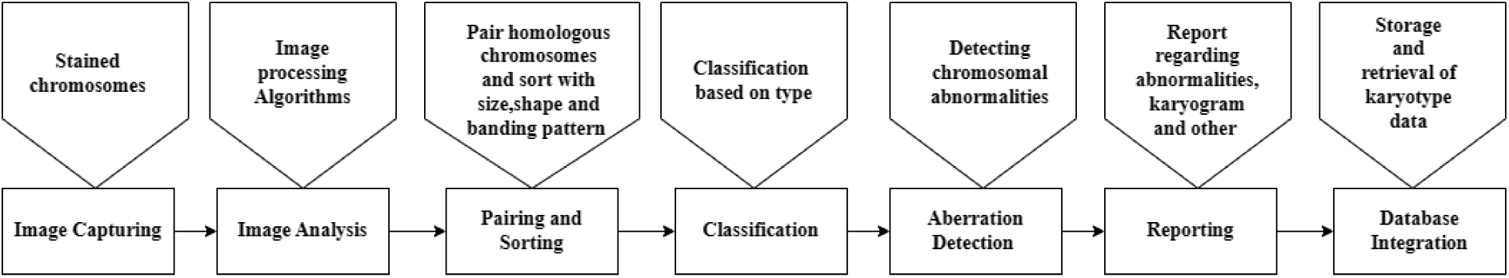
Stages of Computer-Aided Karyotyping System

The computer-aided system will overcome the manual process followed by the cytogeneticists shown in Figure 6. Automated karyotyping faces challenges at every stage, as shown in Figure 7.

**Figure 6:**
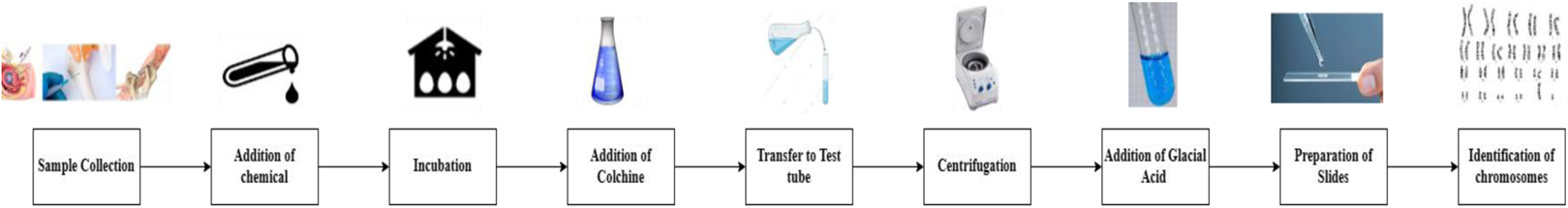
Laboratory process of identification of chromosomes

**Figure 7:**
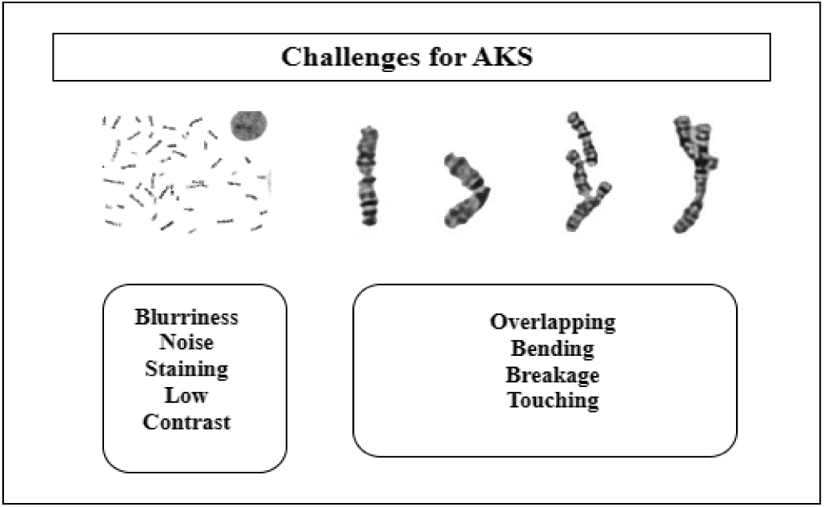
Major Challenges in Automatic Karyotyping System (AKS)

## 2 Research Contributions

The research presented for automating the Karyotyping process has been carried out under this review. The prominent contribution is summarized as follows: -

- The chromosome images follow the staining process to enhance features like band pattern length and centromere position, which are used for pairing and classification.
- The manual karyotyping work has been escalated for automated, efficient classification of genetic diseases in fetuses.
- The presented work has been valid on a standard dataset of Q-banded prometaphase, G-banded, and DAPI images of M-FISH (Multiplex fluorescence in situ hybridization).
- The extracted features face challenges like overlapping, bending, touching, etc.

## 3. Research Methodology

This work followed the Preferred reporting items for systematic reviews and meta-analysis (PRISMA) criteria to conduct an inspection. The chromosomal defects articles in databases, journals, and conferences between 1990 and 2023 have been researched by the review of 150 papers and some of the research because of limited access. After reviewing the articles, the duplicate ones have been omitted from the final list of papers.

The papers from the research publication on Google Scholar, IEEE Xplore, Pub Med, Research Gate, and Science Direct. The free-accessed papers are explored for research. Some papers were free to access and explore. Using the search terms (Machine learning or Deep learning), (Chromosomal disease or Karyotyping or genetic disorder), the papers and articles were chosen. [19–40]. Additionally, articles were chosen based on the exclusion and inclusion criteria. This investigation has been conducted using publications published between 1990 and 2023. After deleting the duplicate papers, only 140 were chosen from 150 research papers. In addition, publications on other diseases were removed from this review article, and the last 130 papers were established and examined with a focus on disorders other than chromosomal conditions.

After that, the entire shortlist of publications was examined, and articles other than genetic algorithms or Machine Learning (ML) or Deep Learning (DL) techniques were deleted. In the end, 123 papers are reported, carefully chosen, and adequately examined. Table 1 will show the PRISMA flow.

**Table 1:**
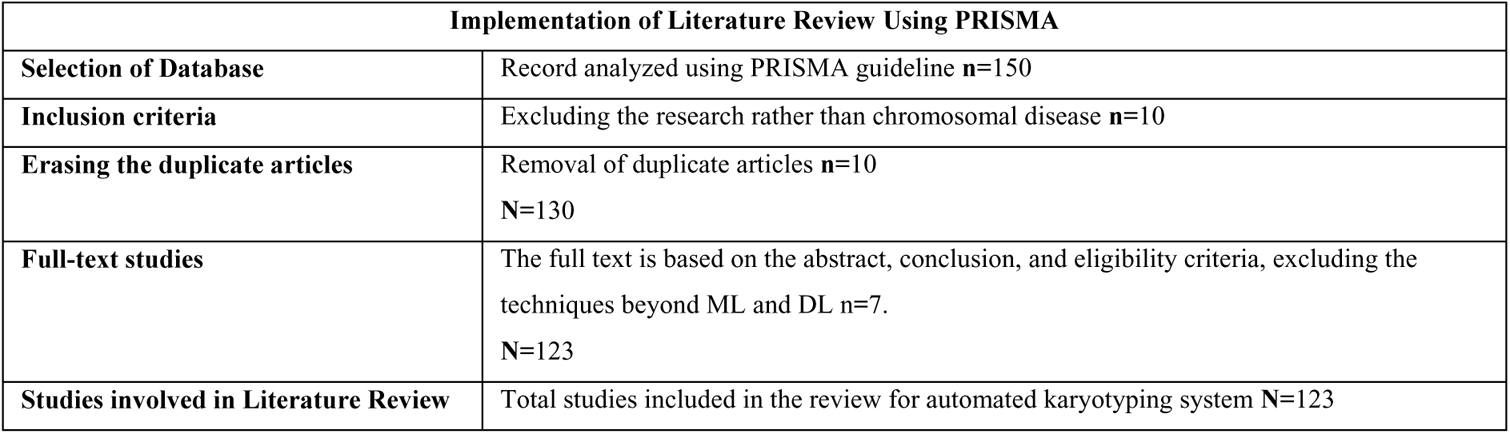
Literature reviewing criteria using PRISMA.

| Implementation of Literature Review Using PRISMA |  |
| --- | --- |
| Selection of Database | Record analyzed using PRISMA guideline n=150 |
| Inclusion criteria | Excluding the research rather than chromosomal disease n=10 |
| Erasing the duplicate articles | Removal of duplicate articles n=10<br>N=130 |
| Full-text studies | The full text is based on the abstract, conclusion, and eligibility criteria, excluding the techniques beyond ML and DL n=7.<br>N=123 |
| Studies involved in Literature Review | Total studies included in the review for automated karyotyping system N=123 |

## 4 Quality Assessment

The inclusion and exclusion criteria were considered while determining the study’s relevance. Most publications cited in the paper offer predictions for various chromosomal disorders based on ML and DL. The paper that has been taken into publication includes empirical research with experimental results.

Additionally, we have compiled a list of the top articles in which scientists utilized ML or DL to forecast a variety of disorders that fall into several categories of chromosomal problems. We compiled the research gleaned from several studies’ abstracts and conclusions for this. The inclusion and exclusion processes followed in the paper are shown in Figure 9.

**Figure 9:**
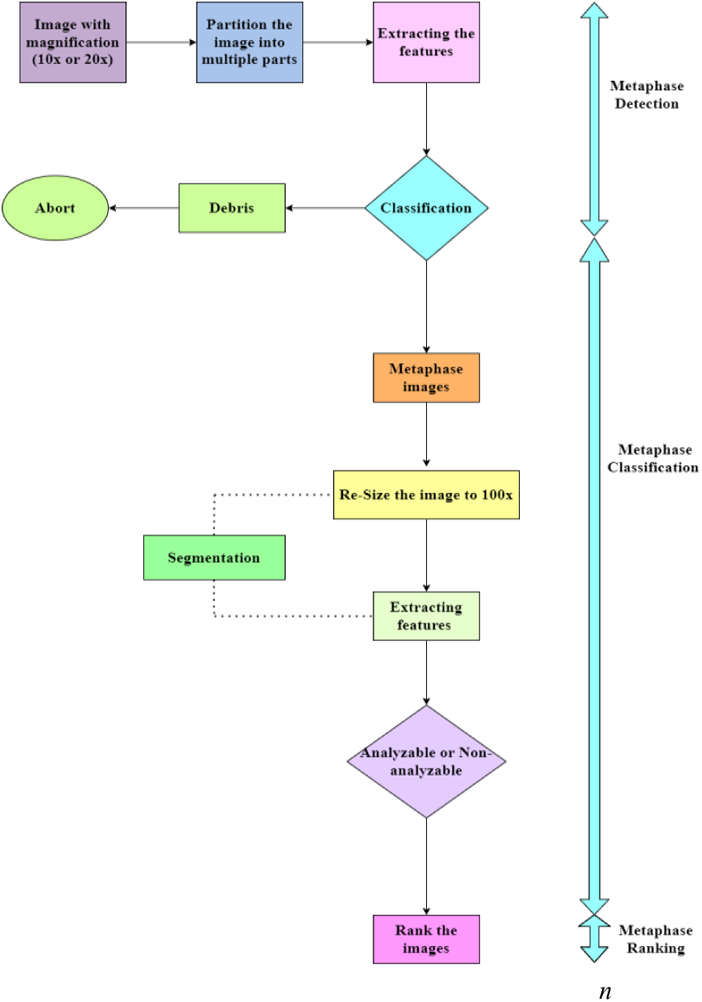
Standards for Inclusion and Exclusion

## 5 Investigations

The review work investigates the following research questions for an automated karyotyping system.

*Investigation 1:* What are the Years-wise publications on Automated Karyotyping Systems?

*Investigation 2:* What are the statistics of publications based on keyword searches in **Google Scholar?**

*Investigation 3:* What are the statistics of Pre-processing techniques used?

*Investigation 4:* What statistics are used to segment chromosomes in automated karyotyping?

*Investigation 5:* What are the statistics of chromosome-used images in automated karyotyping?

*Investigation 6:* What statistic is used to classify chromosomes in automated karyotyping?

*Investigation 7*: What are the statistics of ML and DL techniques used to classify the metaphase images?

## 6 Literature Survey

The automation of the manual process of generating karyogram for the classification of chromosomes and recognizing the abnormalities in chromosomes. The stages followed in the karyotype are Image enhancement, Segmentation, feature extraction, feature selection, and Classification. Every step is vital in generating an automated computer-aided system, as shown in Figure 10.

**Figure 10:**
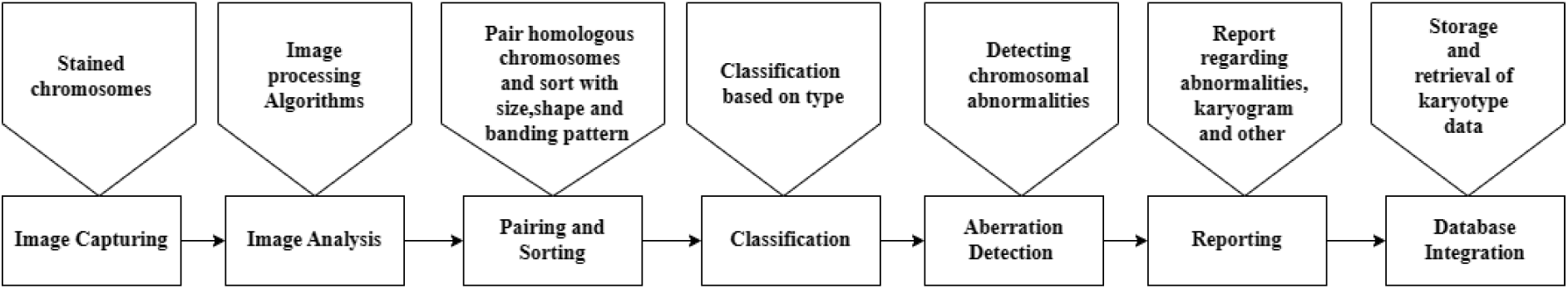
Automated Karyotyping Process (AKS)

### 6.1 Metaphase Selection

The karyotyping system needs to be automated with good-quality metaphase images. A requirement and crucial stage in human karyotyping, which is used to analyze genetic anomalies in people, is the identification of excellent metaphase. The selection of metaphase undergoes the various phases shown in Figure 11. The research based on AKS using different Keywords is shown in Figure 12.

**Figure 11:**
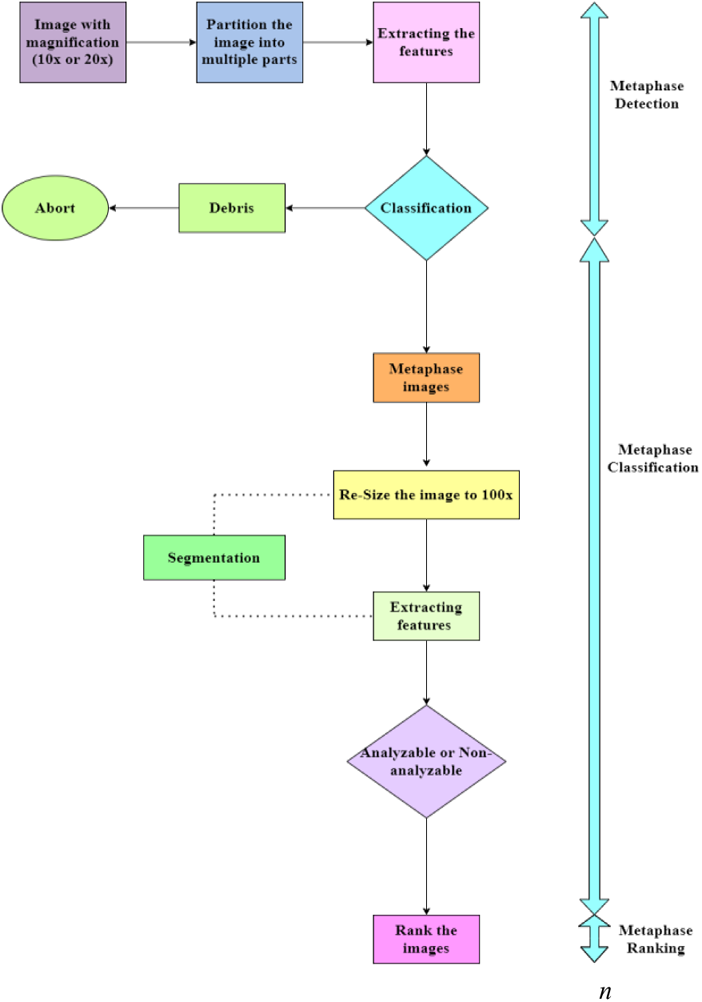
Methodology for metaphase selection

**Figure 12:**
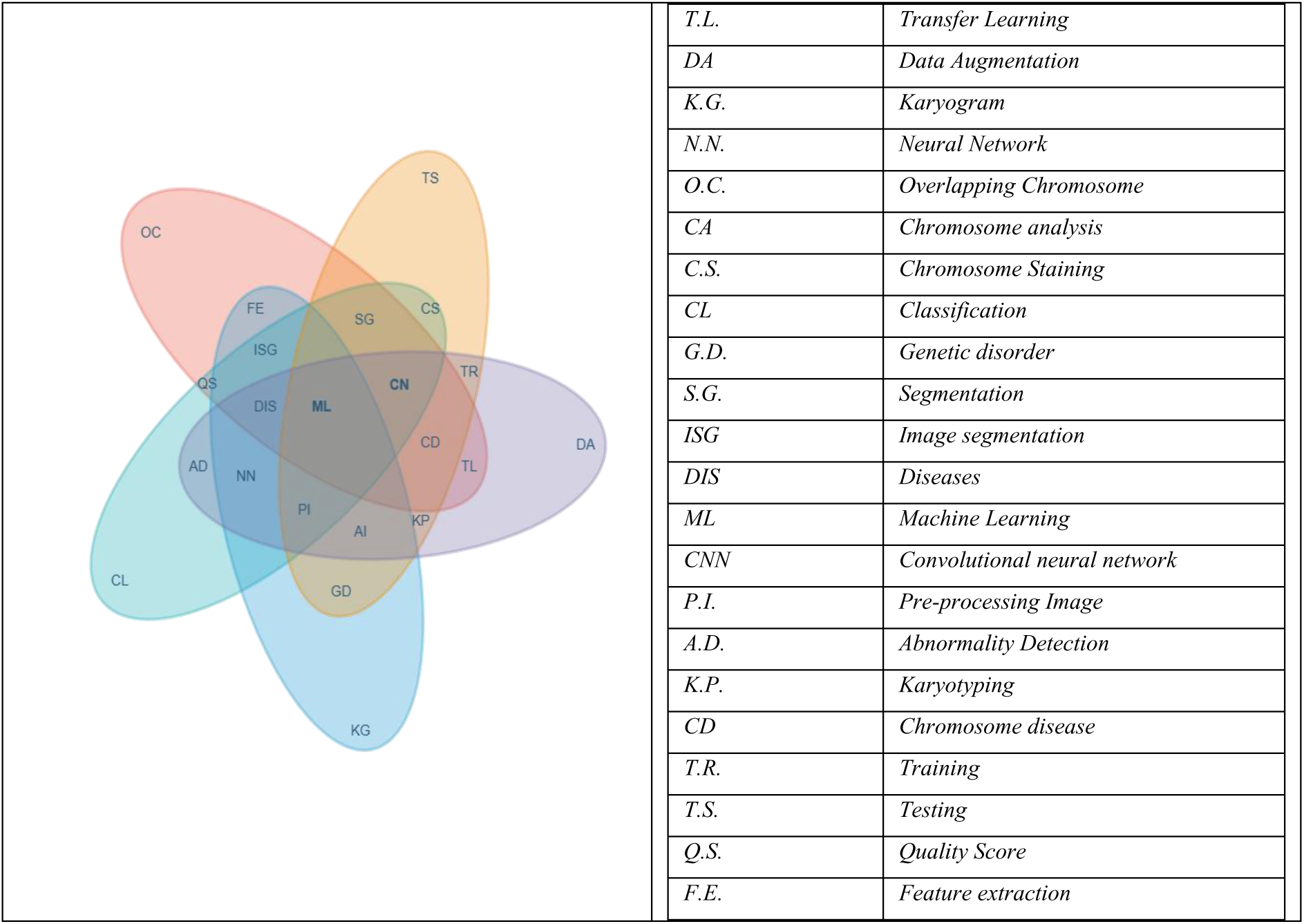
Coincidence of Keywords in Research

The paper demonstrated that METAFER2 would allow an effective computerized identification of meta-phases image selection prepared from mammalian cell lines with different chromosome numbers stained first or division cells. The random metaphase is unaffected by the rising rate of the sister chromatid exchange or various structural chromosomal modifications [7].

The authors work on the automation of mitotic index estimation of cell proliferation and MDWRE (mean depth width ratio of extrema). It also introduced a surface-intensity image roughness characteristic [8].

Chromosome analysis entails discovering good metaphase spreads to identify individuals with genetic issues. However, choosing the right metaphase chromosomes for an accurate clinical assessment takes much time because they are picked manually. The main challenge for conventional cytogenetic analysis is selecting the proper metaphase chromosome spreads. This study uses a software tool based on simple rule-based architecture for karyotyping.[9].

The paper objected to identifying features that allow the proper detection of metaphase chromosomes using high-throughput scanning microscopy. A computer-aided detection (CAD) scheme allows the feature selection for a viable stage because of direct impacts on the scheme’s accuracy [10].

FahamecV1, a low-cost, exact method for metaphase detection, is introduced and discussed in this study. The metaphase stage is when chromosome analysis is done, and the quick and precise automated metaphase stage identification actively shortens the analysis time [11].

The researchers predict the genetic abnormalities by investigating the karyotype in depth. The karyotype is created to organize the chromosomes during the metaphase phase. Chromosomes are flexible bodies that house a person’s genetic material. The findings of segmenting a randomly chosen metaphase chromosomal image might not be accurate and correct [12].

Next, the paper labelled chromosomes using a Deep Assignment Module (DAM). Also, detects the long-range interaction between chromosomes and proposes a Masked Feature Interaction Module (MFIM) that will perform an end-to-end differentiable combinatorial optimization technique known as KaryoNet [13].

The theory of cell proliferation talks about the mitotic index (MI). The MI is often determined by looking at slide preparations under a light microscope. The analyst works on counting a large number of cells and provides the percentage of interphase nuclei with mitotic morphologies [14].

Furthermore, the author selects automated metaphase image needs before the chromosomes are mechanically segmented and categorized. The scholars worked on ranking the meta-spread images(analyzable to least analyzable). Also, count the number of isolated chromosomes, bending, touching, or overlapping chromosomes present in the dataset [15].

The metaphase chromosomal images can be divided into groups that can be analyzed. The approach is more active and more manageable considering geometric considerations for metaphase chromosome image quality. The techniques have been inefficient in comprising images of touching, overlapping and bent chromosomes.

The approach could be more convenient, more active, and more manageable, only considers geometric considerations, has little impact on the quality of metaphase chromosome images, and is ineffective if the image comprises chromosomes in touch, overlap, or bent [16].

A genetic method focused on improving the topology of multi-feature-based artificial neural networks (ANN) and producing images representing all 6900 human chromosomes. The 24 chromosomes were divided into seven categories using one ANN in the first layer of the approach. In the model’s second layer, recognition of seven classes and seven ANNs in specific chromosomes. The method “training-testing-validation” is used for perfect results [17].

A technique known as the MetaSel approach, proposed by the researcher, was used to categorize graded metaphase chromosomal images using eight geometric criteria. Chromosomes that are touching, overlapping or curved can also be treated with it. Its only flaw is the process’s high computing cost of assessing all of the metaphase chromosomal images [18].

However, the paper utilized automated high-throughput scanning microscopy in cytogenetic labs to detect genetic disorders like leukaemia. The primary test uses a unique method to identify analyzable metaphase chromosomes accurately. This project aims to develop a deep learning-based computer-aided detection (CAD) system to precisely identify the chromosomes in metaphase using its eight-layer neural network. The first six layers of the structure are an automatic feature extraction module with a three convolution-max-pooling layer pair architecture. Each first, second, and third pair has thirty, twenty, and twenty feature maps, respectively. Layers seven and eight make up the classifier using multiple-layer perception (MLP) to predict the analyzable metaphase chromosomes. The effectiveness of the new strategy is assessed by utilizing the receiver operating characteristic method. An identification of 150 regions of interest using a novel CAD approach. In the region of interest that contains either interphase cells or metaphase chromosomes [19].

Deep convolutional neural networks are used for the selection stage of a proposed two-step automatic metaphase-finding method and an image processing-based metaphase detection stage. The metaphase images from a 10X scan of the specimen slides. The suggested approach has a 99.33% actual positive rate and a 0.34% false positive rate for metaphase finding [20].

The next researcher uses CNN-LSTM for band sequence property. The input into LSTM is from the CNN-ResNet’s convolutional layers, and the resultant output has been fed into an attention model that assigns 24 labels to its output sequences [21].

In addition, the study proposed a novel method for deep learning-based chromosome defect detection for chromosome feature classification for chromosome abnormality from the dataset with 20,299 chromosome images taken from Dongguan Kanghua Hospital [22]. All approaches are evaluated against a set of parameters in Table 2.

**Table 2:**
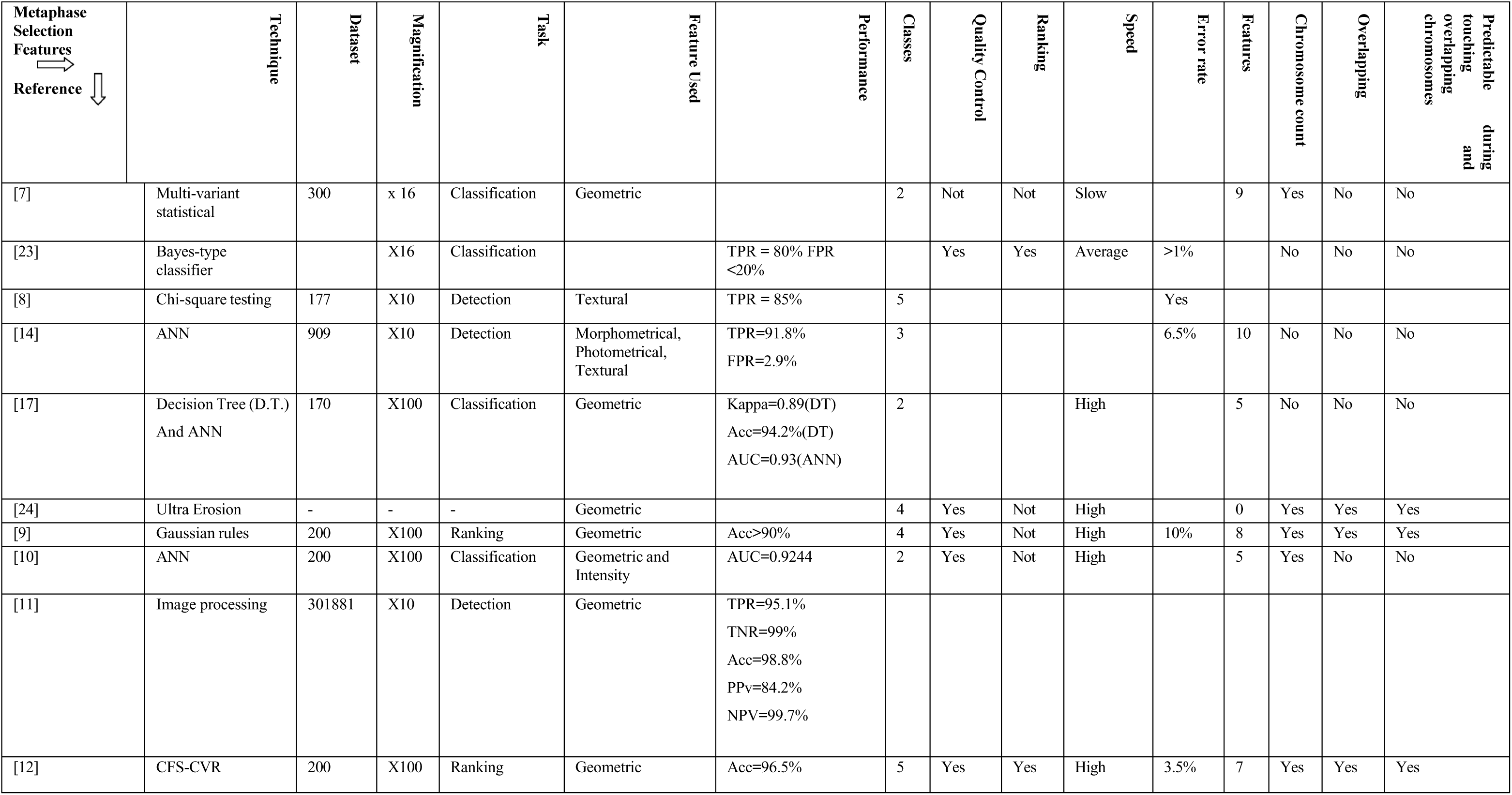
Approached used in Metaphase selection of chromosomes.

| Metaphase Selection Features | Technique | Dataset | Magnification | Task | Feature Used | Performance | Classes | Quality Control | Ranking | Speed | Error rate | Features | Chromosome count | Overlapping | Predictable touching and overlapping chromosomes during |
| --- | --- | --- | --- | --- | --- | --- | --- | --- | --- | --- | --- | --- | --- | --- | --- |
| [7] | Multi-variant statistical | 300 | x 16 | Classification | Geometric |  | 2 | Not | Not | Slow |  | 9 | Yes | No | No |
| [23] | Bayes-type classifier |  | X16 | Classification |  | TPR = 80% FPR <20% |  | Yes | Yes | Average | >1% |  | No | No | No |
| [8] | Chi-square testing | 177 | X10 | Detection | Textural | TPR = 85% | 5 |  |  |  | Yes |  |  |  |  |
| [14] | ANN | 909 | X10 | Detection | Morphometrical, Photometrical, Textural | TPR=91.8%<br>FPR=2.9% | 3 |  |  |  | 6.5% | 10 | No | No | No |
| [17] | Decision Tree (D.T.)<br>And ANN | 170 | X100 | Classification | Geometric | Kappa=0.89(DT)<br>Acc=94.2%(DT)<br>AUC=0.93(ANN) | 2 |  |  | High |  | 5 | No | No | No |
| [24] | Ultra Erosion | - | - | - | Geometric |  | 4 | Yes | Not | High |  | 0 | Yes | Yes | Yes |
| [9] | Gaussian rules | 200 | X100 | Ranking | Geometric | Acc>90% | 4 | Yes | Not | High | 10% | 8 | Yes | Yes | Yes |
| [10] | ANN | 200 | X100 | Classification | Geometric and Intensity | AUC=0.9244 | 2 | Yes | Not | High |  | 5 | Yes | No | No |
| [11] | Image processing | 301881 | X10 | Detection | Geometric | TPR=95.1%<br>TNR=99%<br>Acc=98.8%<br>PPv=84.2%<br>NPV=99.7% |  |  |  |  |  |  |  |  |  |
| [12] | CFS-CVR | 200 | X100 | Ranking | Geometric | Acc=96.5% | 5 | Yes | Yes | High | 3.5% | 7 | Yes | Yes | Yes |

### 6.2 Metaphase picture pre-processing

The chromosome images need to be enhanced for better features like centromere position and length. So, images undergo pre-processing. Staining ineffective samples and imaging conditions are other elements that contribute to decreasing the chromosomal pictures.

The approaches for chromosomal picture enhancement are the main topic of this paper. Investigated are a few algorithms for improvement. These algorithms’ underlying ideas and execution are examined [24].

The pre-processing methods for G-banded metaphase images to create an automated karyotyping system. Photos from the metaphase that can be analyzed are denoised using the median and bilateral filters. Iterative contrast-constrained adaptive histogram equalization enhances denoised images and segments them based on contours [25].

The researchers are visually examining chromosomal banding patterns to identify healthy and abnormal chromosome banding patterns in humans. The changes in chromosomal image display quality, which reduced the cost of the devices, would boost the clarity of the banding pattern, in particular, a detailed discussion of using multiscale transform and derivative filtering for enhancing chromosomal banding patterns [26].

An innovative technique on the cubic-spline wavelet transforms and multiresolution image used for enhancing human chromosomal images. This technique concisely characterizes chromosomal band pattern properties for the transform domain in the multiresolution picture representation. The strategy’s viability and experimental results are given [27].

Chromosome abnormality screening during pregnancy is crucial in ensuring that newborns survive in good health. The pre-processing model with object segmentation and feature improvement has been used. The pre-processing model investigates chromosomal picture feature enhancement and pixel-level chromosome karyotype image extraction [28].

The CIR-contrast improvement ratio, a quantitative metric, is developed to estimate the improved results after comparing an algorithm to numerous existing image-enhancing algorithms. The investigated results predicted from the proposed method are consistently better than the current stage regarding visual effect and the CIR measure [29].

The researchers used a method to improve the contrast of the metaphase images. Exploring the various image enhancement methods like contrast-limiting adaptive histogram equalization(CLAHE), histogram equalization(HE), histogram equalization with 3D block matching(HE+BM3D), and unsharp masking, a more straightforward method. The performance of the contrasted images has been shown with the Structural Similarity Index known as the Peak Signal-to-Noise Ratio (PSNR)[30].

A maximum-likelihood classifier with six features and 25 classes automatically segments and categorizes M-FISH chromosomal pictures. Pre-processing of the photos, background rectification, and the six-channel colour adjustment approach are introduced. Spherical coordinate transformation is a technique for feature transformation. High levels of proper classification are attained [31].

This paper introduces a method known as adaptive and iterative histogram matching for the improvement of the banding pattern of chromosomes. The iteration process produces efficient results with progressive contrast improvement [32].

Enhancing chromosomal contrast using adaptive and iterative histogram matching methods in banding patterns is essential information in chromosome images. The framework of the algorithm consists of computing the statistical moments of the image histogram and establishing each iteration step’s parameters on these moments [33].

The study presented the aim of denoising G-banding of chromosomal images using a cascaded neural network architecture. The process has been covered in two parts. In the first step, a segmentation network was initialized with residual units and U-net abilities. The second stage includes a classification block utilizing the mechanized denoising procedure to eliminate chromosomal loss of pixels [34]. Table 3 explores more literature on the pre-processing study for chromosomes.

**Table 3:** Literature on Pre-processing Techniques.

| Author | Pre-processing Technique | Images Used |
| --- | --- | --- |
| [24] | Sobel operator, high pass filter, low pass filters, Laplacian pyramid, multiscale differential operator. | Digitized chromosome images |
| [26] | Multiscale derivative filtering and three-level steerable pyramid decomposition | Metaphases images |
| [27] | Cubic-spline wavelet transform and multiresolution image analysis | Chromosomes images |
| [29] | Differential wavelet transforms | G-banded chromosome images |
| [31] | Thresholding method | M-FISH Chromosome images |
| [32] | Adaptive and iterative histogram matching (AIHM) | Chromosome images |
| [33] | Adaptive and iterative histogram matching with statistical moments | Not mentioned |
| [34] | Cascaded neural network | G-banding chromosome images |
| [35] | Histogram equalization followed by OSTU thresholding | Chromosome images |
| [36] | Iterative contrast limited adaptive histogram equalization | G banded images |
| [37] | Histogram Equalization with 3D Block Matching (HE+BM3D) | Chromosome images from Meta-System |
| [38] | Contrast Limited AHE (CLAHE) | Chromosome images |

### 6.3 Segmentation

Segmenting chromosome images is necessary before recognizing and identifying certain chromosomal arms. After Segmentation: the objective is to extract the skeleton of the chromosomal arm as a path for a pattern reader to take. The chromosomal bands could be viewed as a form of bar code. There are four subcategories of the widely used chromosomal segmentation algorithm: thresholding, clustering, active contours, and convex-concave point techniques. Segmentation techniques aim to distinguish between chromosomal and background pixels into two groups. The segmentation techniques are independent of object shape; there is little connection between them and the specific problem of chromosomal separation.

As a result, when there is more information, they typically succeed. Since the separation routes of the chromosomes are missing, there is a scenario of incomplete data [39].

Heuristic search edge-linking algorithms seek to separate contacting chromosomes from one another by searching for a least-the cost-connected path that divides the chromosomes. The lost link of such methods to the specific issue of chromosomal separation yields poor results when there is inadequate information, such as when a separating path is absent [40].

A technique that combines the classification step with choosing the best cluster disentanglement to provide Segmentation driven by classification. However, this technique was limited to separating clusters of chromosomes.[41].

With this approach, competing sets of karyotypes are produced. The use of a heuristic system of rules to distinguish between a cluster and a single chromosome, as well as the assumption. The study also points out that, as decided by the segmentation stage, it is tough to deal with images that only contain a small number of single chromosomes or to separate clusters including multiple chromosomes due to the enormous number of variants taken into account. Finally, the various parameters that must be modified and the extensive computation time required raise concerns about the useability in a clinical setting [42].

A method for automatically recognizing, segmenting, and classifying chromosomes from a given metaphase image using a Deep CNN architecture predicts the chromosome type. Chromosome images are segmented, and after that, a Deep CNN model is used to 95.75% accurately classify the images into various classes [43].

Geometric optimizing and multiple input convolutional neural networks (mCNN G.O.) were presented as a method for data classification. After chromosomal straightening with a medial axis locating technique, the segmentation accuracy was 95.644% [44].

The identified overlaps are eliminated by thinning the image using a morphological method. They employed an algorithm to discover the cut sites at the intersection utilizing a given 7×7 mask [46].

The super-resolution layers in the network in the proposed multistage design upscale the image to a higher resolution. The scaled images are subsequently classified using either an Xception or a ResNet50. The best accuracy of 93% was obtained using Xception [47].

To build the SRAS-net classification strategy, the author suggested fusing Deep CNNs with a Super-Resolution Network and a Self-Attention hostile feedback network. They were able to correct the imbalance between the X and Y chromosomal classes with the use of SMOTE to generate extra Y samples. Their accuracy percentage was 97.5% [48].

The author focused on offering sizable segmented datasets that DNN models may use for crowdsourced autonomous categorization. Nevertheless, despite significant automation, this is essentially a human-powered process. The involvement of people in the process is a significant worry [49].

Identifying between partially overlapped chromosomes with a neural network-based picture segmentation technique was possible. On a hypothetical dataset, U-Net segmentation was used to do this. Results [50] showed that with the IoU scores,94.7% were achieved for the overlapped region and 88–94% for the overlapping chromosomal areas. The curvature function scheme, watershed-based Segmentation, region-based, maximum-like hood, pale-path canny edge detection operator, and mathematical morphological operations were some of the different methodologies that the various authors worked on. The literature corresponding to the techniques followed by the researchers is shown in *Table 4*.

**Table 4:** Literature on the Segmentation of metaphase images.

| Author | Year | Approach | Images | Description |
| --- | --- | --- | --- | --- |
| [51] | 2012 | Curvature function scheme | Grey meta spread images | Scheme predict huge number of twisted chromosome structure in the cluster. |
| [52] | 2006 | Watershed based segmentation method | M-FISH | Reduce the computational time |
| [53] | 2008 | Watershed based segmentation method | M-FISH | Calculation of gradient magnitude in Multi Fluorescence In Situ Hybridization images<br>Use of Watershed transform decompose the images into a set of homogeneous regions<br>Achieved at lower computational cost |
| [54] | 1998 | Region based Segmentation | M-FISH | Combinatorial labeling technique<br>Technique with increased the speed and accuracy of image acquisition. |
| [55] | 2001 | Region based Segmentation | G-banding | Multiple thresholds and parameters are heuristically set<br>M-FISH technology's sensitivity and resolution can be increased by using multiplex-labeled region- or locus-specific probes in conjunction with an ideal probe design. |
| [56] | 2004 | Maximum likelihood segmentation | M-FISH | Background correction, color.<br>A compensation and filtering technique as pre-processing step.<br>Lack in the touching or overlapping chromosome. |
| [57] | 2005 | Maximum likelihood hypothesis test | M-FISH | Decomposes both overlap and cluster consisting of more than two chromosomes.<br>Likelihood function has been used to combine chromosome segmentation and classification into a robust chromosome identification system<br>Reliable in indicating errors. |
| [58] | 2009 | Space variant thresholding scheme | Q-banded images | The segmentation method has proven effective even in portions of the image that are highly or poorly luminous.<br>Based on geometric support and picture data, a greedy technique is then utilized to detect and resolve touching and overlapping chromosomes. |
| [59] | 2010 | Pale-path algorithm | Chromosome images | The middle point algorithm extracts the medial axis.<br>The multiscale wavelet method improves the chromosome band. Bi-spline, and extracted using the typical grey, gradient, and shape profiles |
| [60] | 2010 | Watershed based algorithm. | Binary images | Through several pretreatments, overlapping chromosomal pictures were acquired. Then, these photos were processed using the distance transform. The watershed algorithm's input photos were the transformation's results. |
| [61] | 2011 | Modified Snake Algorithm. | M-FISH | In the cluster of metaphase image, touching chromosomes uses a greedy strategy to solve the problem for its separation. |
| [62] | 2012 | Adaptive Fuzzy C-means Clustering Algorithm. | M-FISH | By using a gain field to represent and fix intensity in homogeneities brought on by microscope imaging equipment, flares of targets (chromosomes), and unequal DNA hybridization, the technique enhances the traditional fuzzy c-means algorithm (FCM). |
| [63] | 2012 | Watershed transform | M-FISH | Basic procedures include removing cells from DAPI pictures and gradient calculation, minimum selection, watershed modification, and construction of a binary mask. |
| [64] | 2013 | Canny edge detection operator and mathematical morphological operations. | Gray-scale binary images | The chromosomal are successfully segmented and face problems in the tightly coupled |
| [65] | 2013 | Connected component labelling and extraction. | Q-banded Prometaphase chromosomes | Involves finding of the connected components from the input image and providing labels to produce a Label Matrix for these connected components using the Color Map. |
| [66] | 2013 | Adaptive thresholding method. | G-Band Images | Based on the physical structure of the clusters, there are two basic groups into which clusters fall: single-chromosome and multi-chromosome clusters. |
| [67] | 2014 | MOGAC Algorithm | G-Band Images | A method based on hypothesis curvature is also used to separate and segment chromosomes that have crossed across. |
| [68] | 2014 | Hybrid Technique | G-band Mata spread Images | The four cut spots were utilized to separate the overlapping chromosome images and implementing the linear least squares fit method was employed to generate the chromosome segmentation between point to point. |
| [69] | 2014 | Modified Bacterial Foraging Algorithm | Giemsa-Stained chromosomes images | Bacterium explores the edge pixels in the direction determined by a probabilistic derivative approach derived from ant colony optimization in search of nutrients. |
| [70] | 2015 |  | M-FISH | Removing the various drawbacks of conventional FCM like sensitivity to noise by incorporating both spatial and spectral information. |
| [71] | 2015 | Contour Based | G-Band Metaphase Images | The overlapping zones are obtained together with the chromosomes' outer outline. The curvature function is used for determining the concave and convex points. The overlap zones are located using these concave points. |
| [72] | 2016 | Level Set | ADIR metaphase images. | The Segmentation of images with intensity inhomogeneity can be done pretty effectively using level set-based approaches. |
| [73] | 2016 | Watershed transformation | Giemsa-stained metaphase chromosome images | The identification of centromere candidates, the determination of chromosome width and centerline, and the distinguishing of D.C.s from monocentric chromosomes (MC) using machine learning. |
| [74] | 2017 | Crowdsourcing | Greyscale images | Workers who were partially or not at all marking the chromosomes in their grid vs those who were drawing a broad outline covering every chromosome. |
| [75] | 2017 | Kernel-based adaptive Fuzzy C-Means | M-FISH | Chromosomal preparation using a gain field over a local window for each pixel to account in the intensity in-homogeneities |
| [76] | 2018 | Fully convolutional semantic segmentation network | G-band human chromosomes. | Cross-validation was used to test the semantic segmentation network on samples taken from a public dataset. |
| [77] | 2018 | Binary Watershed Transform | G-band images | The binary mask's distance transform is derived first. Then, grayscale reconstruction is used to process the distance transform negative. |
| [78] | 2019 | Region-Based Active Contours | MFISH | The use of region-based active contour approaches can help overcome the problems brought on by intensity inhomogeneity. These methods find the approximate intensity values along both sides of the contour by using the local intensity values of the adjacent regions of the objects. |
| [79] | 2019 | U-Net-Based Neural Network | Raw G-Band Chromosome Image | The findings of the experiment demonstrated that the convolution neural network may produce satisfying results, particularly with excessively noisy images. |
| [80] | 2020 | CNN model | Microscopic grey images | Addition together the chromosomal overlapping region's pixel values.<br>Different combinations of regions produce a single chromosome |
| [81] | 2020 | Quasi-Newton-based K-means clustering | M-FISH | For the huge and complex issues, the quasi-Newton advantages, including faster computing with balanced results, are ideally suited. |
| [82] | 2021 | Combination of geometry and deep learning | M-FISH | The fully deep learning-based end-to-end chromosomal segmentation techniques rely excessively on pricey labeled data sets. |
| [83] | 2021 | Compact Seg-UNet | DAPI-stained human metaphase chromosomes | To produce more realistic images with opaque overlapping sections to address the issue of unrealistic images in use defined by overlapping regions of higher color intensity. |
| [84] | 2021 | Enhanced Rotated Mask R-CNN | M-FISH | A broad instance segmentation network that may be immediately applied to the karyotyping procedure for chromosomal Segmentation in metaphase images. |
| [85] | 2022 | Adversarial, multiscale feature learning framework | Microscopic grey images | Utilizing open-source datasets for eight evaluation criteria, our framework's performance is demonstrated to be superior, demonstrating its enormous potential for overlapping chromosomal Segmentation. |
| [86] | 2022 | Instance segmentation model based on regression correction | Giemsa-banding chromosome images | The regression-corrected chromosomal instance segmentation network is assessed using an ablation experiment, which demonstrates that the technology can significantly improve performance in automatic chromosome segmentation tasks. |
| [87] | 2022 | Mask R-CNN | Grayscale images | In clusters, the Segmentation of chromosomes with the Mask R-CNN method performed. |
| [88] | 2022 | U-net | G-banding chromosome images | The findings demonstrated that the proposed segmentation network outperforms state-of-the-art semantic segmentation neural networks in terms of dice scores. |

### 6.4 Chromosome classification

Classification has been considered the central part of image analysis. Machine learning provides binary, multi-class and multi-label classification. The dataset undergoes testing, validation and training. The description of a classification process based on local band descriptors. This method’s classification results are contrasted with those using the global band description method (WDD functions) [90].

In a recent study, knowledge-based system techniques were used for the classification of chromosomes, which will overcome the pitfalls of previous methods. The knowledge-based approach has been proposed for the improvement of biomedical patterns. The method used a multistage recursive hypothesize-and-verify paradigm in BPR systems [91].

Additionally, the probabilistic neural network (PNN) technique is used for the classification of chromosomes. The digital dataset was used, and 30 features from each chromosome were extracted. The result consists of 24 separate types of chromosomes, including the 22 autosomes and the X and Y sex chromosomes. Backpropagation has been used for the identification of chromosome classification [92]. *Table 5* shows the research on the classification algorithm.

**Table 5:** Literature on Classification of metaphase images.

| References | Classifier | #Images | Database | Features used | Accuracy % |
| --- | --- | --- | --- | --- | --- |
| [89] | Besiyan | G-banded human metaphase cells. | Dataset 1-Copenhagen (8106 images),<br>Dataset 2-Edinburgh (5469 images)<br>Dataset3-Philadelphia (5817 images) | Band and geometric features | Copenhagen dataset: -91.6%<br>Edinburgh dataset) 80.4 %<br>Philadelphia dataset) 73% |
| [90] | Besiyan | Grey value images | Dataset Copenhagen (7284 images) | Band pattern | 88.5% |
| [91] | Expert system | G-banded human metaphase cell image | Dataset 460 images | Band and geometric features | 91.5% |
| [93] | MLP | G -banding images | Dataset 239 images | Geometrical, band pattern | 76.8% |
| [92] | Transportation Algorithm | - | Dataset 1-Copenhagen (8106 images),<br>Dataset 2-Edinburgh (5548 images)<br>Dataset3-Philadelphia (5947 images) | Band pattern | 95.6% |
| [94] | Compound MLP | Greyscale images | Dataset 1-Copenhagen (8106 images),<br>Dataset 2-Edinburgh (5548 images)<br>Dataset3-Philadelphia (5947 images) | Geometrical, band pattern | Copenhagen dataset: -93.8%<br>Edinburgh dataset) 82.9 %<br>Philadelphia dataset) 77% |
| [95] | MLP | Greyscale images | Dataset 1-Copenhagen (8106 images),<br>Dataset 2-Edinburgh (5469 images)<br>Dataset3-Philadelphia (5817 images) | Band Pattern | Copenhagen dataset: -93.5%<br>Edinburgh dataset) 81.7 %<br>Philadelphia dataset) 77.2% |
| [96] | Probabilistic NN | Grey scale images | Dataset 1-Copenhagen (8106 images),<br>Dataset 2-Edinburgh (5469 images)<br>Dataset3-Philadelphia (5817 images) | Geometrical, band<br>pattern | Copenhagen dataset: -97.0%<br>Edinburgh dataset) 84.7 %<br>Philadelphia dataset) 78.8% |
| [97] | Bayesian |  | Dataset -920 images | Band pattern | 90.32% |
| [98] | MLP | Binary images | Dataset -622 images | Geometrical, density<br>profile | 98% |
| [99] | Hybrid ANN | Grey scale images | Dataset -8818 images | Geometrical, density<br>profile | 66.9% |
| [100] | Non-linear maps |  | Dataset -300 images | Density profile | 86.6% |
| [101] | Multi-layer perceptron<br>(MLP) | Grey level images | Dataset -550 images | Intensity, Geometrical | 83.6% |
| [102] | Backpropagation<br>neural network<br>(BPNN) | Giemsa-Stained images | Dataset -460 images | Geometrical density<br>profile | 93.48% |
| [103] | Fuzzy | Binary images | Dataset-180 images | Geometrical, density<br>profile | 86.7% |
| [104] | Fuzzy |  | Dataset-2400 images | Geometrical, density<br>profile | 96.67% |
| [105] | Gaussian kernel | DAPI image | Dataset-2273 images | Density profile | 93.35% |
| [106] | - | G-banded chromosomes | Dataset-920 images | Geometrical, density<br>profile | 96.89% |
| [107] | Multi-layer perceptron<br>(MLP) | Grey level images | Dataset -Copenhagen(304 images), | - | 88.3% |
| [108] | Heuristic algorithm | Grey level images | Dataset-16094 images | Geometrical, density<br>profile | 94% |
| [109] | Recurrent NN | Grey level images | Dataset-8106 images | Band pattern | 96.1% |
| [110] | ANN | Q-band images | Dataset-8106 images | Density profile and shape | 95.6% |
| [111] | Dynamic time warping | Grey level images | Dataset-1288 images | Geometrical, density profile | 81% |
| [112] | Multi-feature based artificial neural networks (ANN) | Grey level images | Dataset-6900 images | Geometrical, band pattern | 86.7% |
| [113] | Probabilistic ANN | Grey level images | Dataset-2760 images | Geometrical, density profile | 68.19 |
| [114] | Multiscale wavelets Bi-spline | Grey level images | Dataset-lesser than 1000 images | Geometrical, density profile | 85.6 |
| [115] | Wavelet ANN | Grey level images | Dataset-450 images | Geometrical, density profile | 94% |
| [116] | ANN | Q-band images | Dataset-5474 images | Geometrical, density profile | 94% |
| [117] | Multistage Kohonen | Grey level images | - | Geometrical, density profile | 65.5% |
| [118] | MLP | Grey level images | Dataset-5474 images | Geometrical, density profile | 88% |
| [119] | CIR-Net | G-band images | Dataset-2990images | Geometrical | 95.98% |
| [120] | Input-aware and probabilistic prediction convolutional neural network (IAPP-CNN) | Giesma Stain | Dataset-13800 images | Geometrical | 99.2% |
| [121] | Self-Attention Negative Feedback Network (SRAS-net) | Grey scale images | Dataset-5474 images | Geometrical | 97.55% |

## 7 Result and Analysis

### Investigation 1: Years-wise publication on Automated Karyotyping System

The line graph reveals the year-wise publication on Automated Karyotyping System. The trend follows the year 2000 till 2020. There has been an abrupt increase in the research by the researcher. The illustration is shown below in Figure 13.

**Figure 13:**
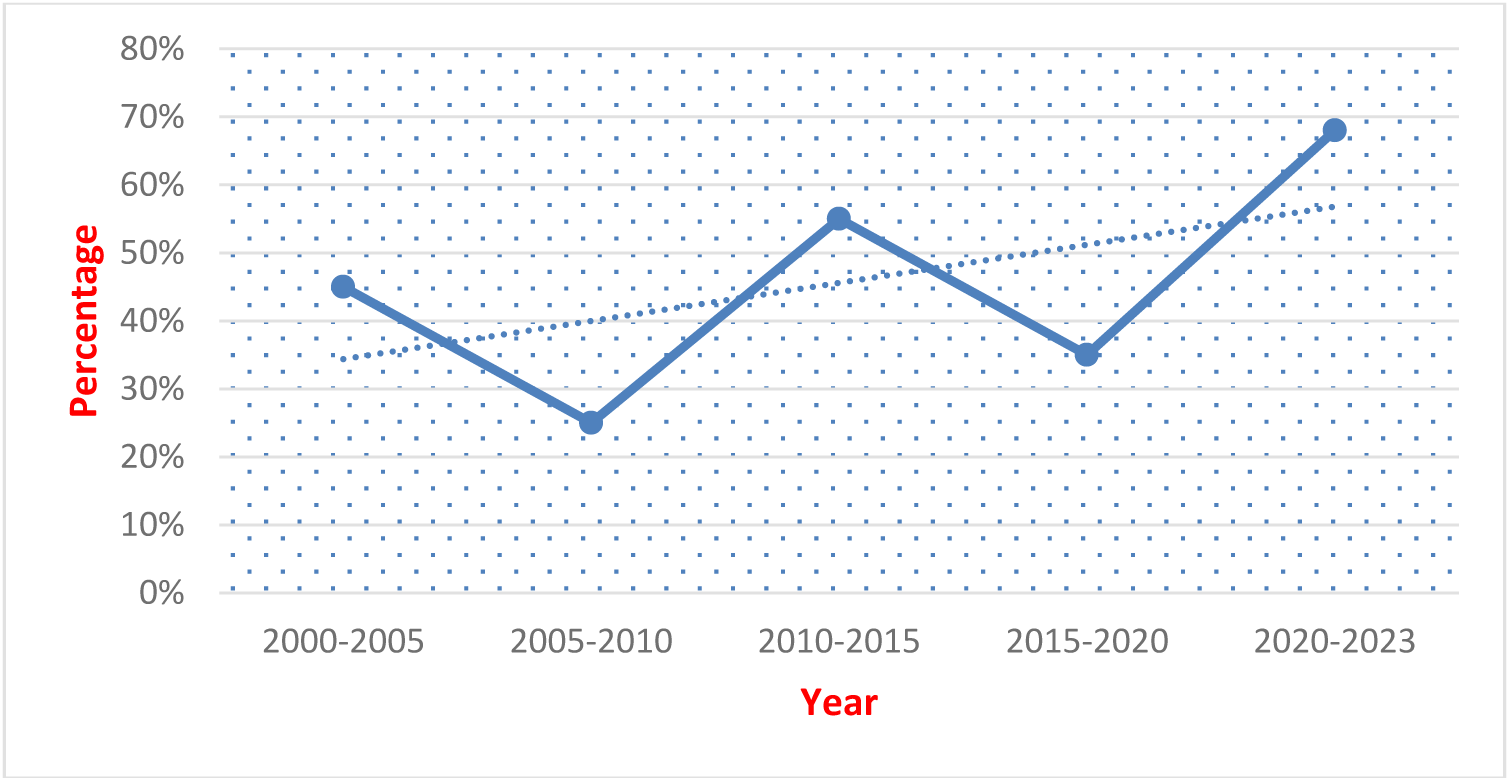
Year-wise research on automated karyotyping system

### Investigation 2: Statistic of publication based on keyword search in Google Scholar

The article published in the area is integrated into the review through keyword research based on the significant repositories. The publications that the demo graphed are excavated from PubMed, IEEE and many more using an advanced keyword search. The graph predicts the year-wise count through the relevant keywords in **Google search** and is shown below in Figure 14.

**Figure 14:**
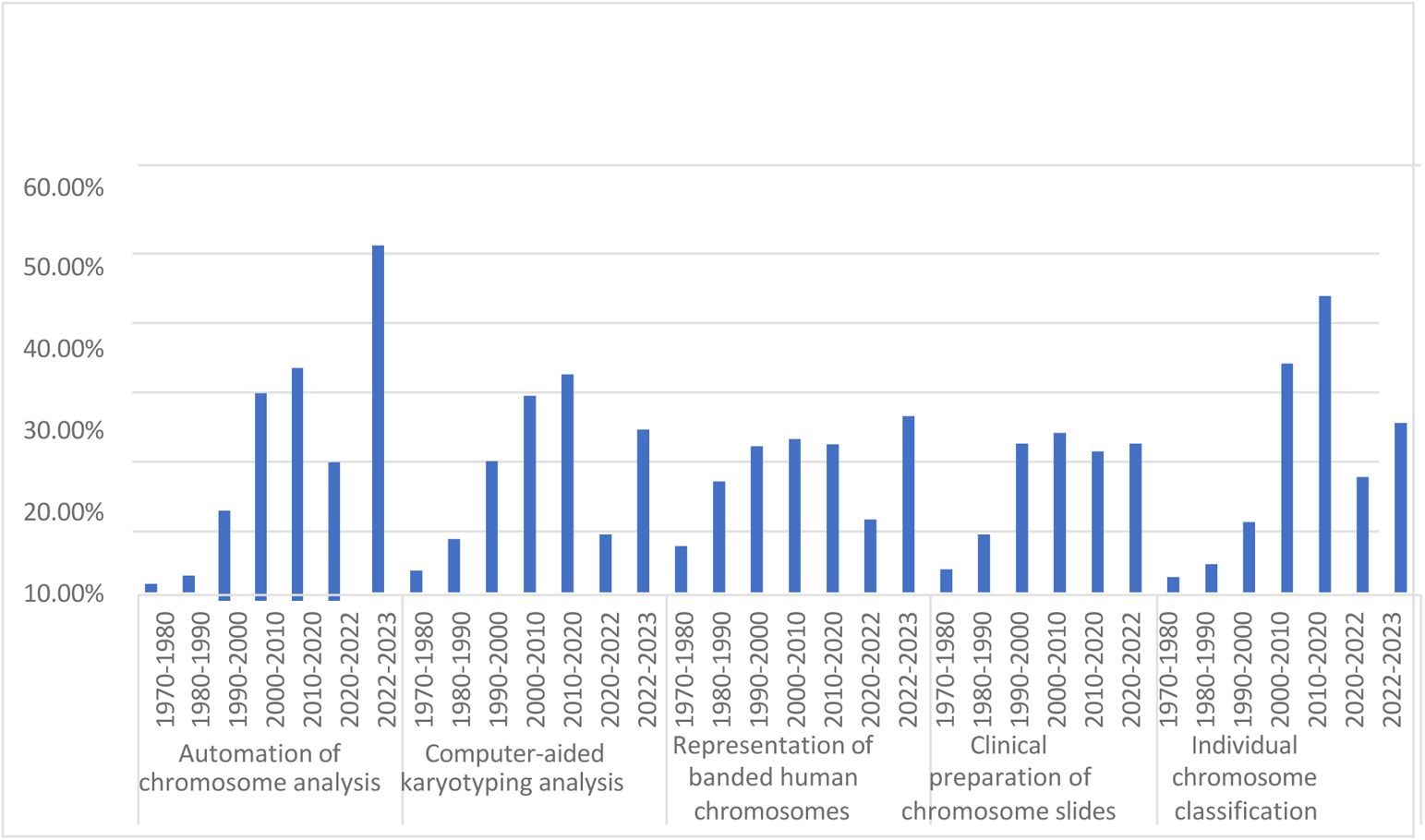
Statistic of publication based on keywords search in Google scholar

### Investigation 3: What are the statistics of Pre-processing techniques used?

Different pre-processing techniques like the Sobel operator, multiscale derivate filtering, Cubic-spline wavelet transform, and many more are used. The methods like cascaded neural networks have been used for pre-processing the metaphase images. The pie-chart depicts the different techniques used in the research of pre-processing of metaphase images shown in Figure 15.

**Figure 15:**
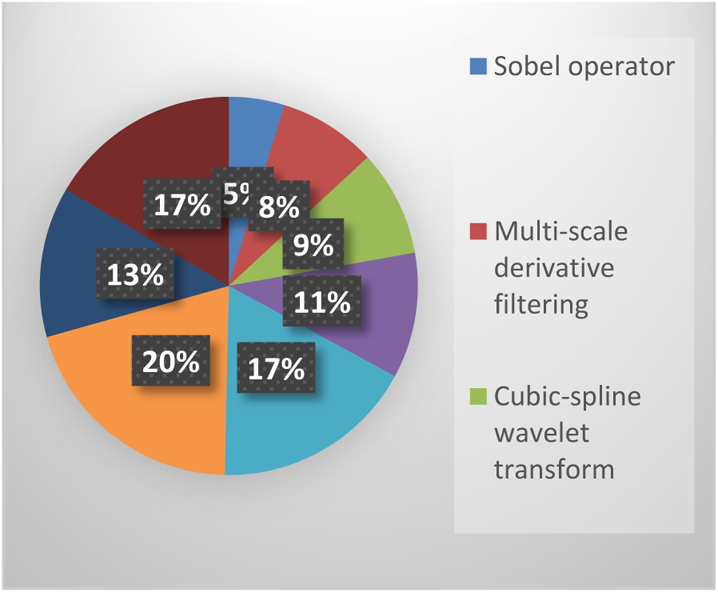
Pre-processing techniques used in research

### Investigation 4: What statistics are used in the Segmentation of chromosomes in automated karyotyping?

The Segmentation of metaphase images is performed using techniques like Watershed, Curvature, Region-based and many more. The pie chart is shown in Figure 16. The year-wise research on the Segmentation is shown in Figure 17.

**Figure 16:**
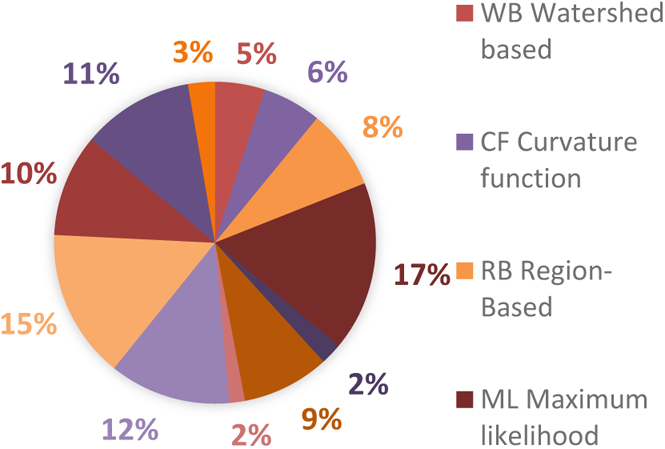
Segmentation techniques

**Figure 17:**
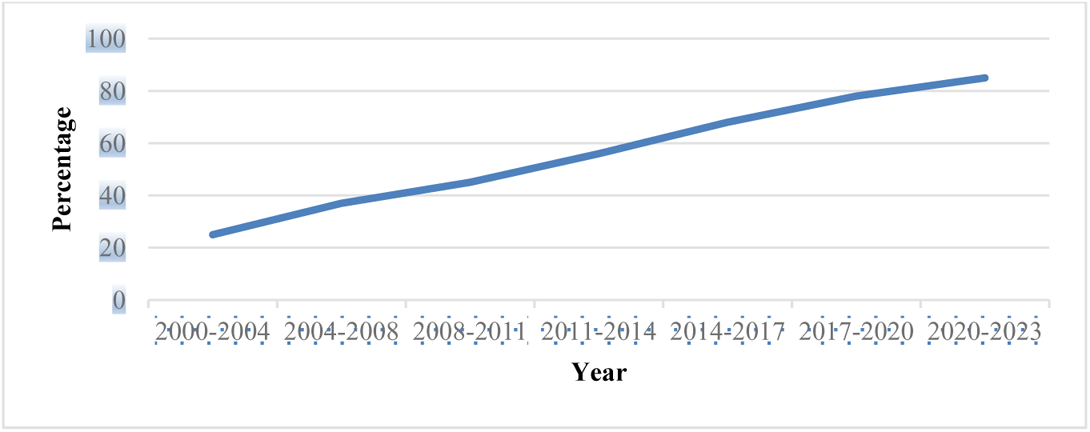
Year wise segmentation of metaphase images of chromosomes

### Investigation 5: What is the statistic of images used in chromosomes in automated karyotyping?

The metaphase images used for classification after staining of the pictures are Giemsa, M-Fish (Multiplex fluorescence in situ hybridization), Q-banded, Greyscale and G-banded. The graph reveals the percentage ratio of images used in automated karyotyping, as shown in Figure 18.

**Figure 18:**
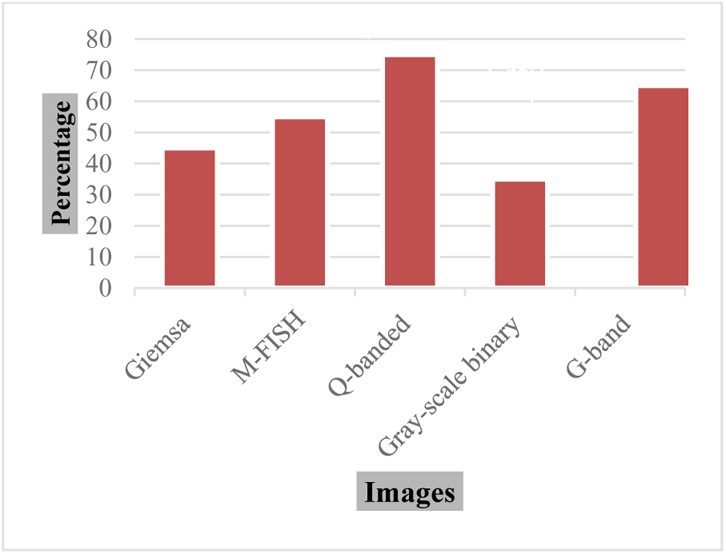
Images used for karyotyping.

### Investigation 6: What statistic is used in the classification of chromosomes in automated karyotyping?

The classification of the extracted features required specified techniques like Besiyan, cascaded neural network, etc. The statistics for type are shown in Figure 19. According to the research, neural-based classification is blooming and helps the authors classify the metaphase images more accurately.

**Figure 19:**
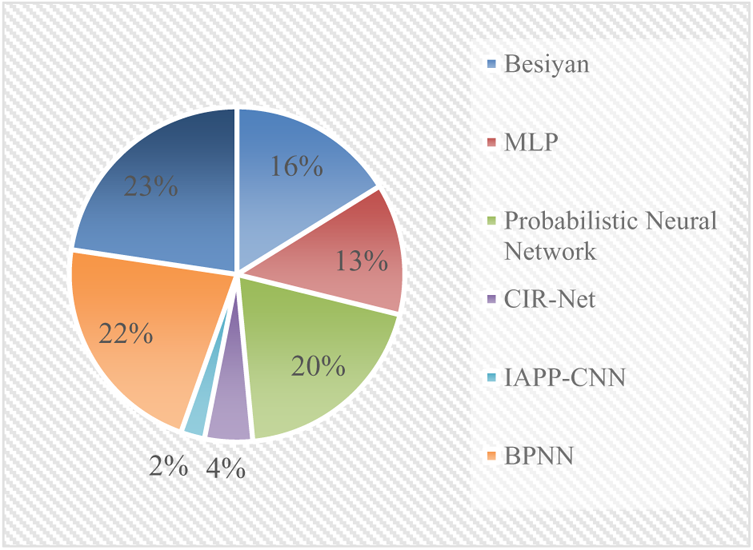
Popular classification techniques

### Investigation 7: What are the statistics of ML and DL techniques used to classify the metaphase images?

The review’s contribution offers concisely updated facts on algorithms developed for the purpose of deep learning and machine learning. The graph explores the algorithm used in the study of the automated karyotyping system, as shown in Figure 20.

**Figure 20:**
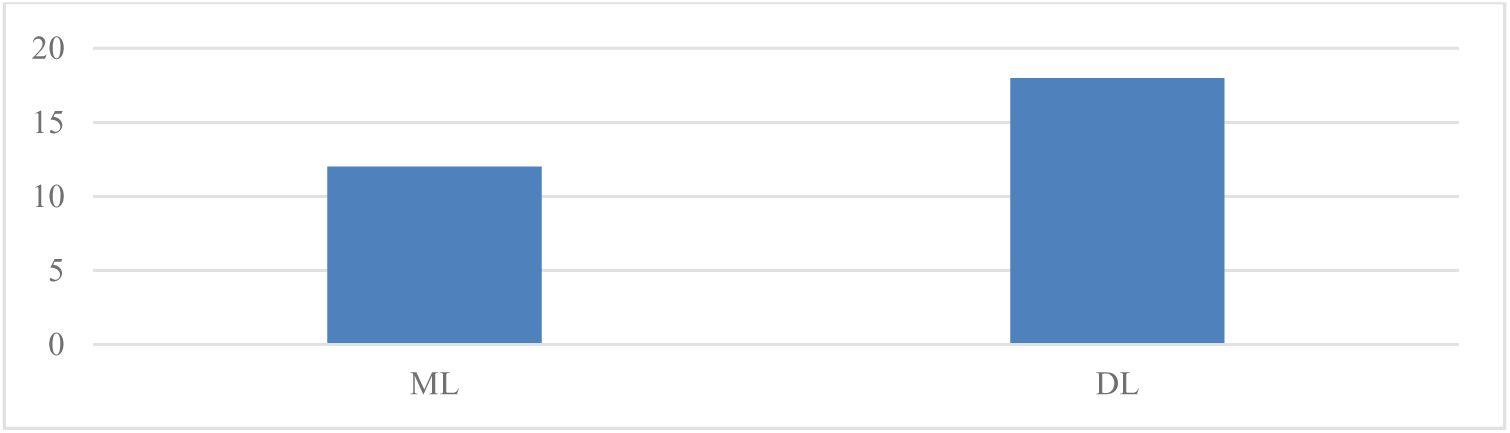
Machine learning (ML) and Deep Learning (DL) techniques used in automated karyotyping system

## 8 Future Direction

This study presents a thorough analysis of many conventional methodologies. This review suggests that future research should move in the following directions. Deep-based approaches can be used to pre-process the automation. Human involvement is still required in karyotyping since touching and overlapping chromosomal separation are crucial. Deep-learning-based segmentation algorithms are solving the rising issue of developing a fully automated system. Deep learning algorithms can be utilized to reconsider chromosomal pairing, even if deep models are broadly used to research classification. Researchers can use the deep models for feature extractors with cutting-edge or custom deep models with or without the involvement of transfer learning. In the latter case, classification can be accomplished using standard ML methods like SVM, random forest, etc. It is possible to create new chromosomal deformation-resistant classification and pairing algorithms.

## 9 Conclusion

This study thoroughly examined several machine learning and deep learning-based methods for computer-aided karyotyping systems to detail genetic illnesses. The limits of several banded pictures, including Q-banded, Gisema, and M-Fish images, were carefully examined. Details are provided regarding the numerous structural and numerical anomalies. We reviewed over 100 articles on mechanical karyotyping systems from reputable journals, including Wiley, Springer, Elsevier, IEEE, PubMed, Taylor and Francis. The data acquired in this study will assist academics in evaluating the most recent advancements in categorizing metaphase photographs. Future research directions must be effectively addressed in light of this substantial literature review. Scheduling technique optimization is a developing research topic. We hope this study’s results will encourage current academics, practitioners, and newcomers interested in novel deep algorithms to address numerous issues noted in the existing literature, thereby advancing the field. It is possible to increase the efficiency of automation technology for karyotyping further by combining various approaches and considering significant critical performance criteria.

